# Vandetanib Reduces Inflammatory Cytokines and Ameliorates COVID-19 in Infected Mice

**DOI:** 10.1101/2021.12.16.472155

**Authors:** Ana C. Puhl, Giovanni F. Gomes, Samara Damasceno, Ethan J. Fritch, James A. Levi, Nicole J. Johnson, Frank Scholle, Lakshmanane Premkumar, Brett L. Hurst, Felipe LeeMontiel, Flavio P. Veras, Sabrina S. Batah, Alexandre T. Fabro, Nathaniel J. Moorman, Boyd L. Yount, Rebekah Dickmander, Ralph Baric, Kenneth H. Pearce, Fernando Q. Cunha, José C. Alves-Filho, Thiago M. Cunha, Sean Ekins

## Abstract

The portfolio of SARS-CoV-2 small molecule drugs is currently limited to a handful that are either approved (remdesivir), emergency approved (dexamethasone, baricitinib) or in advanced clinical trials. We have tested 45 FDA-approved kinase inhibitors *in vitro* against murine hepatitis virus (MHV) as a model of SARS-CoV-2 replication and identified 12 showing inhibition in the delayed brain tumor (DBT) cell line. Vandetanib, which targets the vascular endothelial growth factor receptor (VEGFR), the epidermal growth factor receptor (EGFR), and the RET-tyrosine kinase showed the most promising results on inhibition versus toxic effect on SARS-CoV-2-infected Caco-2 and A549-hACE2 cells (IC_50_ 0.79 μM) while also showing a reduction of > 3 log TCID_50_/mL for HCoV-229E. The *in vivo* efficacy of vandetanib was assessed in a mouse model of SARS-CoV-2 infection and statistically significantly reduced the levels of IL-6, IL-10, TNF-α, and mitigated inflammatory cell infiltrates in the lungs of infected animals but did not reduce viral load.

Vandetanib rescued the decreased IFN-1β caused by SARS-CoV-2 infection in mice to levels similar to that in uninfected animals. Our results indicate that the FDA-approved vandetanib is a potential therapeutic candidate for COVID-19 positioned for follow up in clinical trials either alone or in combination with other drugs to address the cytokine storm associated with this viral infection.

## Introduction

Currently three vaccines are approved for SARS-CoV-2 in the USA ^1, 2, 3^. In contrast, there are relatively few small-molecule drugs that are approved for use including remdesivir ^4^, while molnupiravir is approved in the UK. An emergency use authorization allows the protein kinase inhibitor baricitinib to be combined with remdesivir in hospitalized adults and children 2 years and older who require respiratory support ^5^. The NIH COVID-19 treatment guidelines recommend the use of dexamethasone in certain patients hospitalized with severe COVID-19 based on results from the RECOVERY trial ^6^. As these limited treatment options and several drugs in clinical trials ^7^ attest, there is a global need for more therapeutic options for COVID-19, such as small molecule antivirals that can be used outside of hospitals and treatments to address the many symptoms of COVID-19 that are termed long-COVID ^8, 9^.

Drug repurposing is an essential strategy used to accelerate the discovery of a small molecule treatments for COVID-19 that can enable expedited clinical progression ^10, 11^. Repurposing efforts have already identified molecules (such as remdesivir) originally developed for other viruses and approved outside the USA with potent *in vitro* activity against SARS-CoV-2 ^12^. Small to medium scale assays and high-throughput screening (HTS) campaigns ^13^ have been performed for testing FDA-approved drugs. The catalytic activity of the viral targets M^pro^ and PL^pro^ are essential for viral replication, making inhibition of these enzymes a compelling strategy for antiviral therapy ^14^. The discovery of PF-008352313 as a covalent active-site-directed inhibitor of SARS-CoV M^pro^ in 2003 allowed the translation of this agent into clinical trials for SARS-CoV-2 in 2020 ^15^. Pfizer also developed the SARS-CoV-2 inhibitor PF-07321332 targeting M^pro^, in combination with ritonavir (PAXLOVID^TM^), and an interim analysis of the Phase 2/3 EPIC-HR study showed it reduced risk of death or hospitalization by 89 %. All of these direct acting antivirals target early-stage virus replication and have a short therapeutic window which would render them less effective if administered during the immunopathogenic phase of the disease.

When SARS-CoV-2 invades the body, it can cause an imbalance in the immune system that may result in a cytokine storm ^16^. COVID-19 patients deteriorate over a short period, leading to acute respiratory distress syndrome (ARDS), coagulation disorders, and eventually multiple organ failure ^16, 17^. COVID-19 displays an “inflammatory signature”, characterized by increased levels of soluble biomarkers (cytokine and chemokines) which are involved in the recruitment and activation of several immune cell types, like monocyte/macrophages, neutrophils, T-lymphocytes, and many others ^18^. These immune-active biomarkers, measured either early upon patient admission or throughout hospitalization, may provide clinically relevant information in predicting a more or less severe course of the disease as well as in estimating the mortality risk of infected patients ^19^. Targeting the cytokine storm to ameliorate the state of hyperinflammation has been proposed as a novel therapeutic approach in the treatment of COVID-19 ^18, 20, 21, 22^. It was previously demonstrated for many viruses that the modulation of host cell signaling is crucial for viral replication and it may exhibit strong therapeutic potential ^23, 24^. To date, the FDA has approved over 60 small molecule protein kinase inhibitors and most of these are used in the treatment of cancers (e.g. leukemias, breast and lung cancers) whereas several are for non-malignancies ^25^. Subsequently, protein kinase inhibitors have been proposed for treating SARS-CoV-2 and have demonstrated *in vitro* ^26, 27, 28, 29, 30, 31^ and *in vivo* activity while several are also in clinical trials ^32, 33^. In the current study, we have performed *in vitro* screening of 45 approved protein kinase inhibitors in delayed brain tumor (DBT) cells infected with murine hepatitis virus (MHV), followed by evaluation of entrectinib and vandetanib in A549-ACE2 cells infected by SARS-CoV-2. Finally, we have evaluated the *in vivo* efficacy of vandetanib in an acute infection model using K-18-hACE2 mice challenged with SARS-CoV-2.

## Results

### Vandetanib inhibits SARS-CoV-2 replication *in vitro* without cell toxicity

We used DBT cells infected with murine hepatitis virus (MHV), a model of SARS-CoV-2 replication to evaluate the antiviral activity of 45 kinase inhibitors (Table S1). Among them, 12 showed close to 100% inhibition, and 11 showed moderate activity >60% and <83% at 10 μM (Table S1). Two of the most active hits, entrectinib and vandetanib were also characterized in A549-ACE2 cells infected with SARS-CoV-2 using remdesivir as a positive control. Cells were pretreated with compounds 1 h prior to SARS-CoV-2 infection. Both entrectinib (IC_50_ 1.97 μM) and vandetanib (IC_50_ 0.79 μM) showed lower potency than remdesivir (IC_50_ of 0.11 μM) (Figure 1, A-C). Entrectinib was further tested in Caco-2 and in Calu-3 cell lines infected with SARS-CoV-2, however, this compound was toxic at the concentrations tested (Figure 1D, S1). Entrectinib (5 μM) was tested in Huh-7 cells infected with the human coronavirus 229E (HCoV-229E) ^34, 35, 36^ demonstrating a decrease of 2.7 logTCID50/ml (Figure 1E). In contrast, vandetanib was active in Caco-2 cells (IC_90_ 2 μM, Figure 1D) and showed a reduction of > 3 logTCID50/mL with HCoV-229E when tested at 5 μM (Figure 1 F). Further characterization showed that entrectinib binds to the SARS-CoV-2 Spike RBD with a K_D_ of 166 ± 60 nM (Figure 1G) using microscale thermophoresis (MST) ^37, 38^, an approximately 10 times weaker affinity when compared to ACE2 binding, which has a K_D_ of 15 nM ^39, 40^, which might not be able to protect against virus entry infection and the mechanism might be through inhibition of kinases, since it was reported that growth factor receptor signaling pathways have been reported to be highly activated upon SARS-CoV-2 infection ^41^. SARS-CoV-2 Spike protein-mediated entry was measured in VSV-pseudotype SARS-CoV-2 assays demonstrating that vandetanib was active at 1 μM in the SARS-CoV-2 D614G strain, whereas entrectinib had no significant activity (Figure 1 H, I).

**Figure 1.**
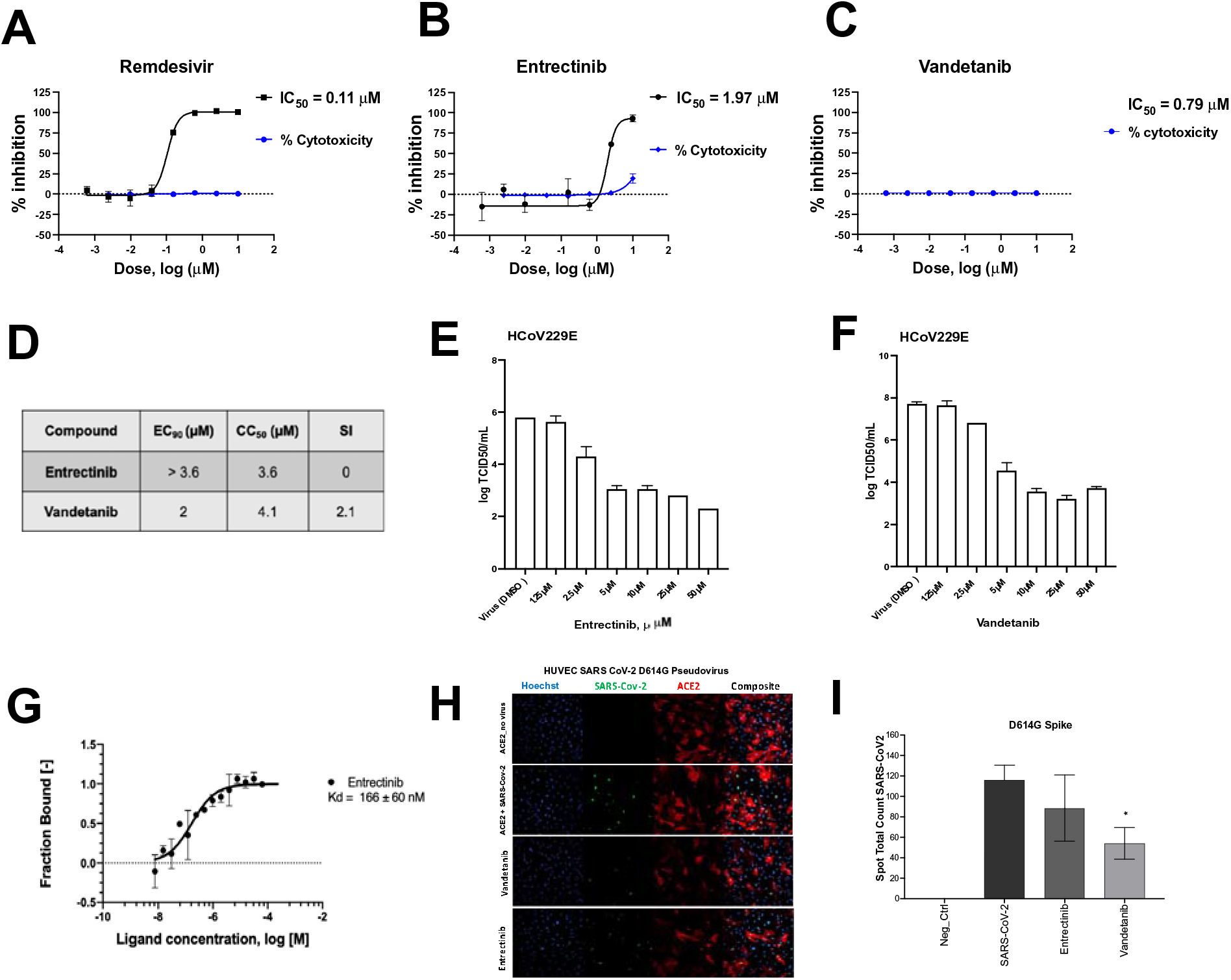
Characterization of entrectinib and vandetanib. SARS-CoV-2 inhibition in A549-ACE2 cell lines and cytotoxicity of **A)** remdesivir **B)** entrectinib and **C)** vandetanib. **D)** EC_90_ and CC_50_ values for entrectinib and vandetanib (strain USA_WA1/2020) in Caco-2 cells. Only VYR data was reported. **E)** HCoV229E antiviral assay and in Huh-7 cell line with entrectinib and **F)** vandetanib. **G)** MicroScale Thermophoresis binding analysis for the interaction between SARS-CoV-2 Spike RBD and entrectinib. **H)** Pseudo SARS-CoV-2 D614G baculovirus (Montana Molecular #C1110G, #C1120G) assay in presence of vandetanib at 1 μM and its **I)** graphical analysis.

### *In vivo* efficacy of vandetanib in mice

Vandetanib *in vivo* efficacy was assessed in the K-18-hACE2 mouse model of COVID-19 ^42, 43, 44^ (8-week-old females, challenged with SARS-Cov-2 2 x10^4^ PFU, intranasally). Vandetanib (25 mg/kg) was administered i.p. 1 h before infection and once daily up to day 3 post-infection (3 dpi) (Figure 2A). On 3 dpi, mice were euthanized, and viral load, cytokines and lung histopathology were evaluated. In all groups tested, mice lost weight compared to uninfected animals that received only vehicle formulation (Figure 2B). Lung viral load was evaluated by RT-PCR, and, although vandetanib reduced SARS-Cov-2 infection in A549-ACE2 cells, no significant reduction in viral RNA levels was observed *in vivo* when compared with the infected untreated mice (Figure 2C). While vandetanib did not decrease the viral load, it had a clear protective effect on lung pathology (Figure 2D and E). Infected untreated mice showed severe pathological changes with inflammatory cell infiltrates. In contrast, vandetanib treatment exhibited improved morphology and milder infiltration even in the absence of effect on viral load. These results might indicate an effect on the virus-induced inflammatory process. Further analyses also showed that vandetanib treatment restored the levels of IFN-1β (Figure 3A and B) and prevented the increase of the levels of the most widely evaluated inflammatory cytokines/chemokines observed in infected mice. Vandetanib reduced IL-6, TNF-α, and CCL4 (compared to infected untreated animals) to similar levels found in uninfected animals (Figure 3C, D and E). Vandetanib also significantly reduced the levels of CCL2, CCL3, and IL-10 compared to infected animals (Figure 3F, G and H). CXCL1 was not affected by the treatment (Figure 3I). CXCL2 and CXCL10 were not elevated in infected animals (Figure 3J and K). Altogether, these results indicate that the *in vivo* efficacy of vandetanib in the COVID-19 mouse model is more likely related to the reduction in the cytokine storm than reduction in SARS-CoV-2 replication.

**Figure 2:**
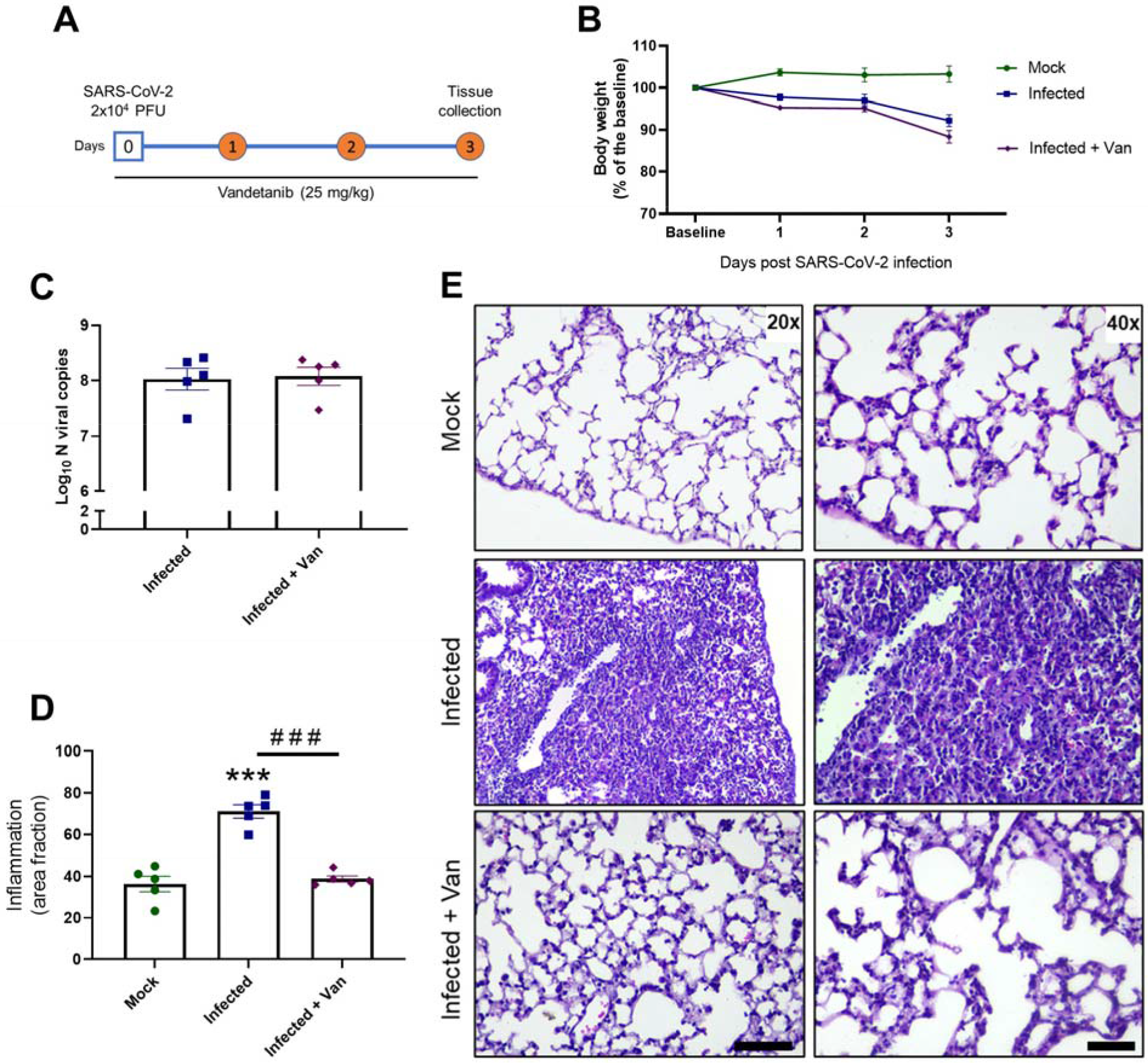
*In vivo* efficacy of Vandetanib in a mouse model of COVID-19. A) Experimental timeline: K18-hAce2 tg mice were infected with SARS-CoV-2 (2 × 10^4^ PFU/40 μL saline, intranasal) or mock. One group of mice was treated with Vandetanib (25 mg/kg i.p.) 1 h before virus inoculation. B) Body weight was evaluated daily. C) At 3 DPI, mice were euthanized and the C) lung viral load, and D-E) Lung histopathology were evaluated. *** p<0.001 as compared with mock group after one-way ANOVA followed by Tukey post-hoc test. ### p<0.001 as compared with infected group after one-way ANOVA followed by Tukey post-hoc test. Bar scales = 20x – 125 μm; 40x – 50 μm.

**Figure 3:**
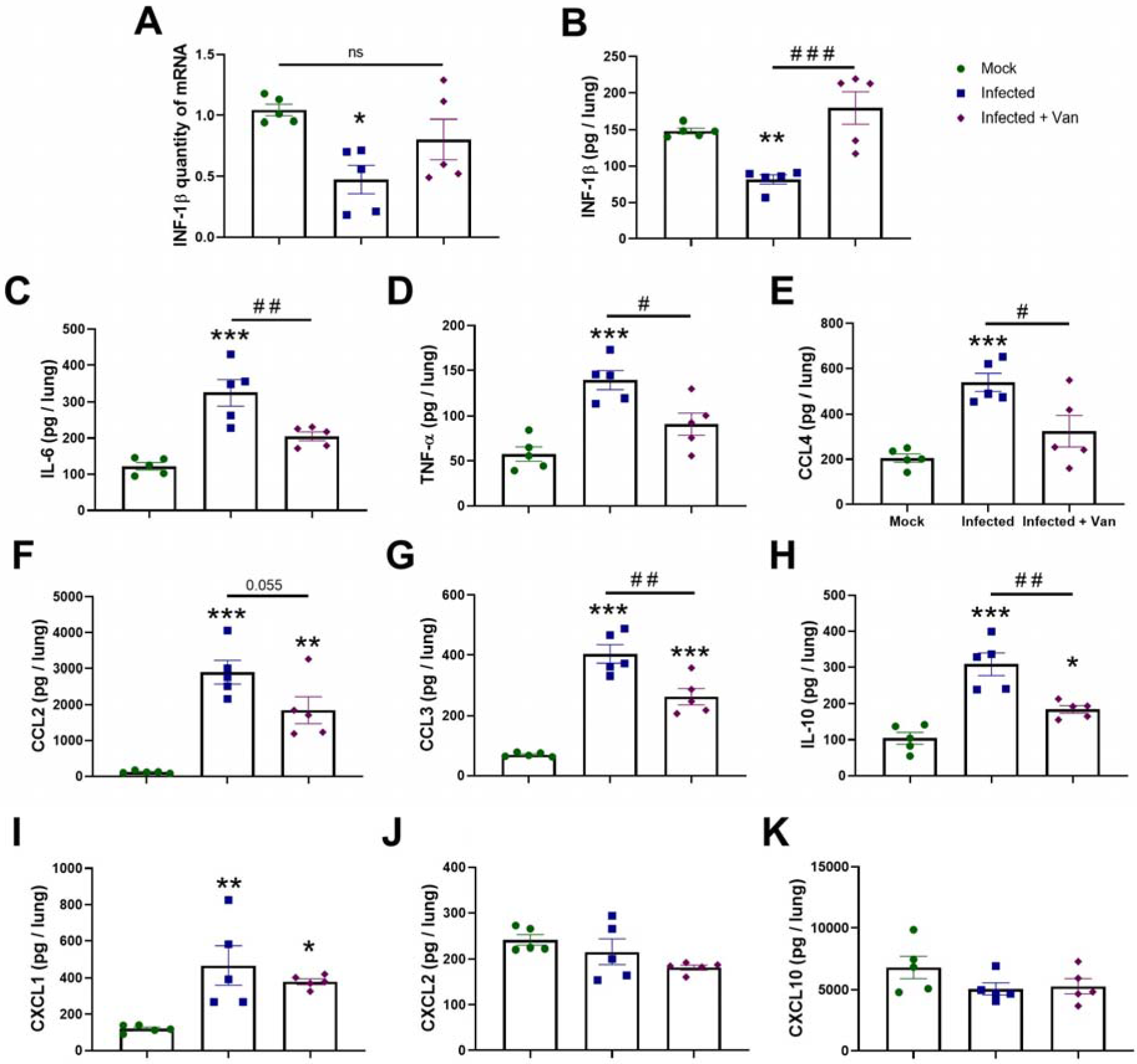
Effects of Vandetanib on SARS-CoV-2-induced lung inflammation in a mouse model of COVID-19. A) Expression of INF-1β quantified by qPCR. Levels of B) INF-1β, C) IL-6, D) TNF-α, E) CCL4, F) CCL2, G) CCL3, H) IL-10, I) CXCL1, J) CXCL2, and K) CXCL10 measured by ELISA. * p<0.05, ** p<0.01, and *** p<0.001 as compared with mock group after one-way ANOVA followed by Tukey post-hoc test. # p<0.05 and ## p<0.01 as compared with infected group after one-way ANOVA followed by Tukey post-hoc test.

## Discussion

With a mounting infection and death toll to date, there has been a focus on ensuring the global population are vaccinated against SARS-CoV-2. Still, we are in a race against rapidly emerging viral variants that may hamper the vaccines effectiveness in future ^45^, while many countries still have little or no access to the vaccines that are available in the USA and Europe ^46^. There is also a critical need to develop treatments that do not require cold chain storage and can be used outside of hospitals. Repurposing approved small molecule drugs represents the fastest route to the clinic if they can demonstrate statistically significant efficacy in animal models.

Growth factor receptor signaling pathways have been reported to be highly activated upon SARS-CoV-2 infection, hence inhibition of these pathways prevents replication in cells ^47^. After screening 45 kinase inhibitors against MHV, we narrowed them down to entrectinib and vandetanib. Vandetanib is an FDA approved drug used to treat thyroid gland tumors (targeting VEGFR, EGFR and RET-tyrosine kinase ^48^) and was active in Caco-2 cell, A549-ACE2 infected by SARS-CoV-2 and against HCoV-229E. Based on this *in vitro* activity profile it was selected for testing in a COVID-19 mouse model.

One of the major causes of ARDS and multi-organ dysfunction syndrome (MODS) observed in severe SARS-CoV-2 infection is the cytokine storm ^17, 49^. Studies have revealed higher levels of cytokine storm associated with more severe COVID development ^17^. In these patients, accumulation and exudation of inflammatory substances from the cytokine storm destroys tissues, leading to multi-organ failure and ARDS ^49^, an important cause of death in severe and critical cases of COVID-19. Capillary leakage is also major component of deteriorating lung function in COVID-19, resulting in ARDS due to inflammation driven by TNF, IL-1, IL-6, IL-8, and VEGF ^17^. Abnormal coagulation can result in organ failure and death in patients with severe COVID-19, caused by the cytokine storm ^17^. IL-1, IL-6, TNF, signal transducer and transcriptional activator 3 (STAT3), NF-kB, and lipopolysaccharides were all identified as regulators of thrombotic markers in COVID-19 patients with the severe-to-critical disease ^50^. We are not aware of any current therapies that target the cytokine storm that have reversed disease.

Our evaluation of the *in vivo* efficacy of vandetanib in a murine model of infection demonstrated that vandetanib reduced IL-6, IL-10, and TNF-α compared to infected untreated animals to similar levels found in uninfected animals. In patients with severe COVID-19, the levels of several inflammatory cytokines and chemokines including PDGF, TNF, IL-6, and VEGF are significantly increased ^51^. IL-6 has also been shown to correlate with respiratory failure and adverse clinical outcomes ^49, 52^. Furthermore, a recent clinical study showed that the ARDS group had higher levels of IL-6, IL-8, and IL-10 than the non-ARDS group, and the levels of these cytokines correlated significantly with coagulation parameters and disseminated intravascular coagulation (DIC) ^49^. The levels of IL-6 and TNF-α correlated with the levels of creatinine and urea nitrogen and were also higher in ARDS patients with acute kidney injury (AKI) ^49^. Thus, reducing the levels of these mediators upon treatment with vandetanib could improve prognosis in COVID-19. The host innate immune response releases cytokines such as type I interferon (IFN α and β) during infection, thereby initiating antiviral activity However, this particular interferon response is interrupted by factors such as SARS-CoV-2 non-structural proteins, aging, diabetes, and germ-line errors eventually making the host more susceptible to illness ^53, 54, 55, 56, 57^. One interesting observation is that vandetanib rescued the decreased IFN-1β caused by SARS-CoV-2 infection in mice to levels similar to that in uninfected animals (Figure 3A). Vandetanib also decreased CCL2, CCL3, CCL4 and CXCL1 compared to infected animals (Figures 3F, 3G, 3E, 3I). These chemokines have been reported to be increased in patients with COVID-19 ^19^. CXCL1 is a known chemoattractant for neutrophils, and it was identified, among others, to be the one of the more abundantly expressed by macrophages involved in SARS-CoV2 infection, especially in more severe cases ^58^ CCL2 is one of the key chemokines that regulate migration and infiltration of monocytes/macrophages and CCL2 levels were also independently associated (together with other immune soluble biomarkers) with mortality in COVID-19 patients ^59^. Finally, RNA isolated from nasopharyngeal swab samples from COVID-19 positive patients showed that CCL2 (and CCL3) expression was significantly higher in patients with unfavorable outcomes than cases with a favorable evolution or with negative controls ^60^. Targeting chemokine/chemokine receptor binding might suppress hyper immune activation in critical COVID-19 patients^58^.

The lung histological examination for mice treated with vandetanib showed a significant treatment effect with mild infiltration, looking similar to uninfected mice. ARDS and dyspnea create hypoxia in lung tissues and other organs. Hypoxia induces VEGF expression. VEGF is a potent vascular permeability factor that induces vascular leakiness in COVID-19 infected lung tissues, resulting in plasma extravasation and pulmonary edema, which further increases tissue hypoxia ^61, 62^ and VEGF participates in lung inflammation ^63^. Blocking VEGF and the VEGF receptor (VEGFR)-mediated signaling would improve oxygen perfusion and anti-inflammatory response and alleviate clinical symptoms in patients with severe COVID-19. In agreement, bevacizumab, a humanized anti-VEGF monoclonal antibody was employed for treating patients with severe COVID-19 (NCT04275414 ^64^). Relative to comparable controls, bevacizumab showed clinical efficacy by improving oxygenation and shortening oxygen-support duration, and by day 28, 92% of patients show improvement in oxygen-support status, 65% patients were discharged, and none showed worsened oxygen-support status nor died ^64^. Therefore modulating VEGF using drugs such as vandetanib ^48^ may have clinical utility.

Many FDA-approved kinase inhibitors have been proposed as broad-spectrum antiviral therapies ^65^ because they have multiple protein targets, including those in the host cell required for viral life cycle, replication, and infection of multiple virus types. Kinase inhibitors also have anti-inflammatory and cytokine inhibitory activity properties which may address lung damage from respiratory virus infections ^65^. Host-target antivirals also offer the advantage that they can exploit the host protein and pathways needed for the replication of the virus and resistance may be less likely to develop against them.

Although we observed vandetanib reduced infection in i) DBT cells infected by MHV and in ii) A549-ACE2 infected by SARS-CoV-2, iii) showed a reduction of > 3 logTCID50/mL of HCoV-229E and iv) decreased viral entry in the pseudovirus assay, we did not observe a statistically significant reduction in the viral load in the murine SARS-CoV-2 infected model. We would likely need much longer *in vivo* studies to demonstrate a reduction in viral replication. We also did not observe a decrease in remdesivir viral load although it has been previously reported that remdesivir (25mg/kg) subcutaneous studies in mice should use *Ces1c^-/-^* which lack the serum esterase in order to mirror pharmacokinetics exposure that is seen in humans ^66^. While our results may represent a suboptimal remdesivir dose it demonstrated a positive effect on the lung pathology in mice comparable to vandetanib. The positive effects shown in our study regarding modulation of the main inflammatory cytokines and chemokines as well as prevention of lung damage and fibrosis, demonstrated that vandetanib likely has the potential to address the cytokine storm associated with SARS-CoV-2 infection. Although the antiviral remdesivir was shown to reduce the length of hospital stay for those with COVID-19 ^67^, only anti-inflammatory approaches have improved survival in these patients, such as dexamethasone when given to those with an oxygen requirement ^68^. Most recently, a randomized, placebo-controlled trial of Janus-kinase inhibition using tofacitinib has been reported to improve COVID-19 survival, in the presence of background glucocorticoid treatment (received by 89% of patients) ^69^. The SAVE-MORE trial showed that blockade of the cytokine IL-1 via the recombinant human IL-1 receptor antagonist anakinra in patients with COVID-19, guided by suPAR levels in patients hospitalized with moderate and severe COVID-19 significantly reduced the risk of worse clinical outcome at day 28 ^70^.

Minimizing or preventing the cytokine storm is still a significant challenge, as it will require definition of the timing immunosuppressive or immunomodulatory agent administration. Knowing the cytokines to target in severe COVID-19 pneumonia and providing more-targeted therapeutic approaches may allow for the earlier introduction of anti-cytokine treatment. We now report that vandetanib can decrease levels of several such important cytokines that are significantly elevated in the cytokine storm, such as IL-6 and TNF-α.

In conclusion, there is continued interest in kinase inhibitors for treating COVID-19 and several such as masitinib ^71^ and others are in clinical trials ^26^. We now report that the FDA approved kinase inhibitor vandetanib could be a potential drug to target the cytokine storm and prevent patients from developing ARDS. Treatment with vandetanib in the mouse model also reduced key inflammatory cytokines. Vandetanib is well absorbed from the gut, reaching peak blood plasma concentrations 4 to 10 h after application, and has an average half-life of 19 days ^72, 73^. The pharmacokinetic properties of vandetanib were linear over the dosage range of 50 to 1200 mg/d. The maximum plasma concentration of 857 ng/mL is usually reached after 6 h in patients with thyroid medullary cancer at a dose of 300 mg. Vandetanib is highly protein-bound (92–94%) and has a terminal excretion half-life of 20 d. It is metabolized by cytochrome P450 3A4 (CYP3A4) and is predominantly excreted via the feces and urine (44% and 25%, respectively ^48^). In another study in healthy patients with dose escalation up to 1200 mg/d, vandetanib appeared to be well tolerated in the populations studied and at the dose of 800 mg/kg, vandetanib can reach a C_max_ of 1 μM ^72^, that is above the IC_50_ reported in our study infection of SARS-CoV-2 in A549-ACE2 cells. When combined these pharmacokinetic effects and positive impacts on the cytokine storm and preventing lung inflammation, may suggest it is worthy of clinical studies for COVID-19. While ee are aware of at least one report of a patient treated with vandetanib who had COVID-19 and recovered ^74^, to date it has not been accessed further in a clinical trial which this current study may point towards.

## Acknowledgments

We graciously thank Dr. Sara Cherry and Dr. David Schultz for the Calu-3 high-content SARS-CoV-2 studies performed by the University of Pennsylvania High-throughput Screening Core and the Cherry laboratory. Dr. Mindy Davis and colleagues are gratefully acknowledged for assistance with the NIAID virus screening capabilities. The authors gratefully acknowledge the technical assistance of Marcella Daruge Grando, Ieda Regina dos Santos, Juliana Trench Abumansur, and Felipe Souza. We kindly thank Dr. Anne. M. Quinn from Montana Molecular for her generous assistance with pseudovirus testing.

## Grant information

We kindly acknowledge NIH funding: 1R43AT010585-01 from NIH/NCCAM to SE, AI142759 and AI108197 to RSB. This project was also supported by the North Carolina Policy Collaboratory at the University of North Carolina at Chapel Hill with funding from the North Carolina Coronavirus Relief Fund established and appropriated by the North Carolina General Assembly.

J.A. Levi and N.J. Johnson were supported by the Comparative Medicine Institute at North Carolina State University through its CAVE initiative.

Dr. Glaucius Oliva and colleagues acknowledge Coordenação de Aperfeiçoamento de Pessoal de Nível Superior (CAPES – project 88887.516153/2020-00) and Fundação de Amparo à Pesquisa do Estado de São Paulo (FAPESP project 2013/07600-3).

Collaborations Pharmaceuticals, Inc. has utilized the non-clinical and pre-clinical services program offered by the National Institute of Allergy and Infectious Diseases. TMC, JCAF and FQC received funding from the São Paulo Research Foundation (FAPESP) under grant agreement 2013/08216-2 (Center for Research in Inflammatory Disease) and 2020/04860-8 and from Coordenação de Aperfeiçoamento de Pessoal de Nível Superior (project 88887.507155/2020-00).

Collaborations Pharmaceuticals, Inc. has utilized the non-clinical and pre-clinical services program offered by the National Institute of Allergy and Infectious Diseases.

## Supplemental Materials

### METHODS

#### Chemicals and reagents

Entrectinib, Vandetanib were purchased from MedChemExpress (MCE, Monmouth Junction, NJ).

#### Expression and purification of Spike RBD of SARS-CoV-2

A codon-optimized gene encoding for SARS-CoV-2 (331 to 528 amino acids, QIS60558.1) was expressed in Expi293 cells (Thermo Fisher Scientific) with human serum albumin secretion signal sequence and fusion tags (6xHistidine tag, Halo tag, and TwinStrep tag) as described before ^1^. S1 RBD was purified from the culture supernatant by nickel–nitrilotriacetic acid agarose (Qiagen), and purity was confirmed to by >95% as judged by coomassie stained SDS-PAGE. The purified RBD protein was buffer exchanged to 1x PBS prior to analysis by Microscale Thermophoresis.

#### Microscale Thermophoresis

We used Microscale thermophoresis (MST) to detect binding of entrectinib to the Spike RBD protein. The experiments were performed according to the manufacturer’s instructions (NanoTemper) and as described previously ^2^. Briefly, for protein labeling, 6 μM of protein was be used with 3-fold excess NHS dye in MST Buffer (HEPES 10 mM pH 7.4, NaCl 150 mM), using Monolith Protein Labeling Kit RED-NHS 2nd Generation (Amine Reactive). Free dye was removed, and protein eluted in MST buffer, and centrifuged at 15 k rcf for 10 min. Binding affinity measurements were determined using NanoTemper’s Monolith NT.115 Pico (Nanotemper) and were performed using 5 nM protein a serial dilution of compounds, starting at 100 μM in MST buffer containing 5 % glycerol, 1 mM β-Mercaptoethanol and 0.1 % Triton X-100. Spike RBD was incubated at room temperature in presence of compounds for 20 min prior measurement. Samples were then loaded into sixteen standard capillaries (NanoTemper Technologies) and fluorescence was recorded for 20 s using 20 % laser power and 40 % MST power. The temperature of the instrument was set to 23°C for all measurements. After recording the MST time traces, data were analysed. KD value was calculated from ligand concentration-dependent changes in the fraction bound (Fbound) of Dye-Spike RBD after 10 s of thermophoresis. The assay was performed in quadruplicate and the values reported were generated through the usage of MO Affinity Analysis software (NanoTemper Technologies).

#### Pseudovirus Assay

Cell imaging and analysis was conducted at Phenovista Biosciences. HUVEC single cell donor (Lonza, cat#C2517A) cells were transduced at room temperature with ACE2 using a BacMam viral vector at a concentration of 2e^9^ VG/ml (Montana Molecular #C1120G Pseudo SARS-CoV-2 D614G Green Reporter) followed by incubation at 36°C for 24 hours. After this step, inhibitor compounds were diluted to 1μM and incubated for 60 minutes with 2e9 VG/ml of Pseudo SARS-CoV2 or Pseudo SARS-CoV2 D614G baculovirus (Montana Molecular #C1110G, #C1120G). Prior to fixation with PFA, cell nuclei were stained with Hoechst, and images were acquired with the high content screening InCell Analyzer HS6500 microscope (20X Magnification). Quantitative analysis was done with ThermoFisher HCS Studio Cell analysis suite.

#### SARS-Cov-2 tested in A549-ACE2 cells

A549-ACE2 cells were plated in Corning black walled clear bottom 96 well plates 24 hours before infection for confluency. Drug stocks were diluted in DMSO for a 200X concentration in an 8-point 1:4 dilution series. Prepared 200X dilutions were then diluted to 2X concentration in infection media (Gibco DMEM supplemented with 5% HyClone FetalCloneII, 1% Gibco NEAA, 1% Gibco Pen-Strep). Growth media was removed, and cells were pretreated with 2 X drug for 1 hour prior to infection at 37C and 5% CO2. Cells were either infected at a MOI of 0.02 with infectious clone SARS-CoV-2-nLuc ^3^ or mock infected with infection media to evaluate toxicity. 48 hours post infection wells were treated with Nano-Glo Luciferase assay activity to measure viral growth or CytoTox-Glo Cytotoxicity assay to evaluate toxicity of drug treatments, performed per manufacturer instructions (Promega). Nano-Glo assays were read using a Molecular Devices SpectraMax plate reader and CytoTox-Glo assays were read using a Promega GloMax plate reader. Vehicle treated wells on each plate were used to normalize replication and toxicity. Drug treatment was performed in technical duplicate and biological triplicate.

#### SARS-Cov-2 tested in Calu-3 cells

Calu-3 (ATCC, HTB-55) cells were pretreated with test compounds for 2 hours prior to continuous infection with SARS-CoV-2 (isolate USA WA1/2020) at a MOI=0.5. Forty-eight hours post-infection, cells were fixed, immunostained, and imaged by automated microscopy for infection (dsRNA+ cells/total cell number) and cell number. Sample well data was normalized to aggregated DMSO control wells and plotted versus drug concentration to determine the IC_50_ (infection: blue) and CC_50_ (toxicity: green).

#### SARS-Cov-2 tested in Caco-2 cells

For the Caco-2 VYR assay, the methodology is identical to the Vero 76 cell assay other than the insufficient CPE is observed on Caco-2 cells to allow EC50 calculations. Supernatant from the Caco-2 cells are collected on day 3 post-infection and titrated on Vero 76 cells for virus titer as before.

#### Murine Hepatitis Virus

Each compound was tested for antiviral activity against murine hepatitis virus (MHV), a group 2a betacoronavirus, in DBT cells. MHV-A59 with nano-Luciferase: The MHV-A59 G plasmid was engineered to replace most of the coding sequence for orf4a and 4b with nano-luciferase (nLuc). Briefly, nucleotides 27,983 to 28,267 were removed and replaced with SalI and SacII restriction sites; approximately 111 bp of the 3’ end of orf4B was left to maintain the TRS for orf5. Nano-luciferase was pcr amplified with primers 5’nLuc SalI (5’-NNNNNNGTCGACATGGTCTTCACACTCGAAGATTTC-3’) and 3’nLuc SacII (5’-NNNNNNCCGCGGTTACGCCAGAATGCGTTCGCAC-3’), digested with SalI and SacII and then cloned into the G plasmid which had been similarly digested. A sequence verified G-nLuc plasmid was used with MHV-A59 wild type A, B, C, D, E and F plasmids to recover virus expressing nLuc, using our previously described molecule clone (Systematic assembly of a full-length infectious cDNA of mouse hepatitis virus strain A59 ^4^. Each compound was tested against MHV using an 8-point dose response curve consisting of serial fourfold dilutions, starting at 10 μM. The same range of compound concentrations was also tested for cytotoxicity in uninfected cells.

#### HCoV 229E antiviral assay

HCoV 229E, (a gift from Ralph Baric, UNC, Chapel Hill) was propagated on Huh-7 cells and titers were determined by TCID_50_ assay on Huh-7 cells. Huh-7 cells were plated at a density of 25,000 cells per well in 96 well plates and incubated for 24 h at 37°C and 5% CO_2_. Growth media was removed, and cells were pretreated with 2 X drug for 1 hour prior to infection at 37C and 5% CO2. Cells were infected with HCoV 229E at a MOI of 0.1 in a volume of 50 ul MEM 1+1+1 (Modified Eagles Medium, 1% FBS, 1% antibiotics, 1% HEPES buffer) for 1 hour. Virus was removed, cells rinsed once with PBS growth medium was added back at a volume of 100 μl. Supernatants were harvested after 24 h, serially ten-fold diluted, and virus titer was determined by TCID_50_ assay on Huh-7 cells. CPE was monitored by visual inspection at 96h post infection. TCID_50_ titers were calculated using the Spearmann-Karber method ^5, 6^.

#### Mouse studies

##### Ethical approval

All the experimental procedures were performed in accordance with the guide for the use of laboratory animals of the University of Sao Paulo and approved by the institutional ethics committee under the protocol number 105/2021.

#### SARS-CoV-2

SARS-CoV-2 was isolated from a COVID-19 positive-tested patient. The virus was propagated and titrated in Vero E6 cells in a biosafety level 3 laboratory (BSL3) at the Ribeirao Preto Medical school (Ribeirao Preto, Brazil). Cells were cultured in DMEM medium supplemented with 10% fetal bovine serum (FBS) and antibiotic/antimycotic (Penicillin 10,000 U/mL; Streptomycin10,000 μ Vero cells in DMEM (FBS 2%) incubated at 37 °C with 5% CO_2_ for 48 h. The cytopathogenic effect was observed under a microscope. Cell monolayer was collected, and the supernatant was stored in −70 °C. Virus titration was made by the plaque-forming units (PFU).

#### K18-hACE2 mice

To evaluate the effects of vandetanib *in vivo*, we infected the K18-hACE2 humanized mice (B6.Cg-Tg(K18-ACE2)2Prlmn/J)^7, 8, 9^. K18-hACE2 mice were obtained from The Jackson Laboratory and were breed in the Centro de Criação de Animais Especiais (Ribeirão Preto Medical School/University of São Paulo). This mouse has been used as model for SARS-CoV-2-induced disease and it presents clinical signs, and biochemical and histopathological changes compatible with the human disease^8, 9, 10, 11, 12, 13, 14^. Mice had access to water and food *ad libitum*. For the experimental infection, animals were transferred to the BSL2 facility.

#### SARS-CoV-2 experimental infection and treatments

Female K18-hACE2 mice, aged 8 weeks, were infected with 2×10^4^ PFU of SARS-CoV-2 (in 40 μL) by intranasal route. Uninfected mice were inoculated with equal volume of PBS. On the day of infection, 1 h before virus inoculation, animals were treated with vandetanib (25 mg/kg, i.p.) (n = 6). Five infected animals remained untreated. Vandetanib was also given once daily on the days 1, 2 and 3 post-infection. Body weight was evaluated on the baseline and on all the days post-infection. On the day 3 post-infection, 6 h after treatments, animals were humanely euthanized, and lungs were collected. Right lung was collected, harvested, and homogenized in PBS with steel glass beads. The homogenate was added to TRIzol reagent (1:1), for posterior viral titration via RT-qPCR, or to lysis buffer (1:1), for ELISA assay, and stored at −70 °C. The left lung was collected in paraformaldehyde (PFA 4%) for posterior histological assessment.

#### Absolute viral copies quantification

Total RNA from the lung was obtained using the Trizol® (Invitrogen, CA, EUA) method and quantified using NanoDrop One/Onec (ThermoFisher Scientific, USA). A total of 800 ng of RNA was used to synthesize cDNA. cDNA was synthesized using the High-Capacity cDNA Reverse Transcription kit (Applied Biosystems, Foster City, CA, USA), following the manufacturer’s protocol. The determination of the absolute number of viral copies was made by a taqman real-time qPCR assay with the ad of the StepOneTM Real-Time PCR System (Applied Biosystems, Foster City, CA, USA). A standard curve was generated in order to obtain the exact number of copies in the tested sample. The standard curve was performed using an amplicon containing 944 bp cloned from a plasmid (PTZ57R/T CloneJetTM Cloning Kit Thermo Fisher®), starting in the nucleotide 14 of the gene N. To quantify the number of copies, a serial dilution of the plasmid in the proportion of 1:10 was performed. Commercial primers and probes for the N1 gene and RNAse P (endogenous control) were used for the quantification (2019-nCov CDC EUA Kit, IDT), following the CDC’s instructions.

#### ELISA assay

Lung homogenate was added to RIPA buffer in proportion of 1:1, and then centrifuged at 10,000 g at 4 °C for 10 minutes. Supernatant was collected and stored in −70 °C until use. The Sandwich ELISA method was performed to detect the concentration cytokines and chemokines using kits from R&D Systems (DuoSet), according to the manufacturer. The following targets were evaluated: IL-6, IL-10, IL-1β, TNF-α, INF-1β, CCL2, CCL3, CCL4, CXCL1, CXCL2, and CXCL10.

#### Lung histopathological process and analyses

Five micrometer lung slices were submitted to Hematoxylin and Eosin staining. A total of 10 photomicrographs in 40X magnification per animal were randomly obtained using a microscope Novel (Novel L3000 LED, China) coupled to a HDI camera for images capture. The total septal area and total area were analyzed with the aid of the Pro Plus 7 software (Media Cybernetics, Inc., MD, USA). Morphometric analysis was performed in accordance with the protocol established by the American Thoracic Society and European Thoracic Society (ATS/ERS) ^15^.

**Figure S1:**
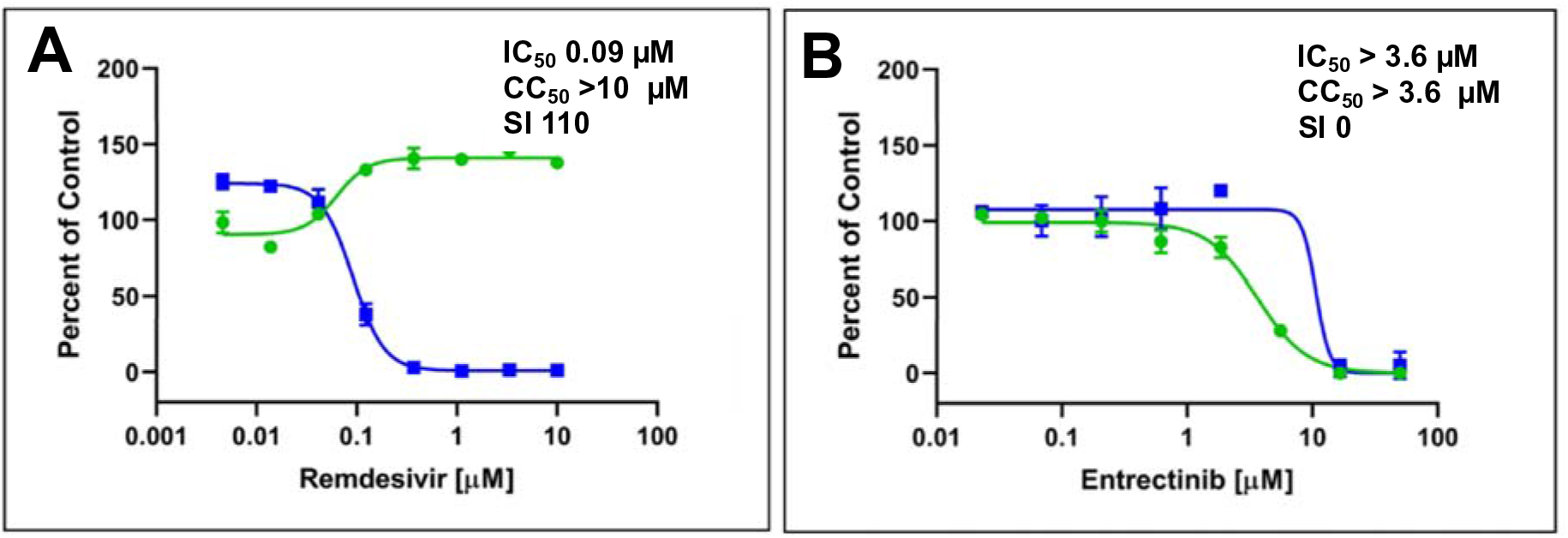
*In vitro* antiviral SARS-CoV-2 testing in Calu-3 cells. Calu-3 (ATCC, HTB-55) cells were pretreated with test compounds for 2 hours prior to continuous infection with SARS-CoV-2 (isolate USA WA1/2020) at a MOI=0.5. Forty-eight hours post-infection, cells were fixed, immunostained, and imaged by automated microscopy for infection (dsRNA+ cells/total cell number) and cell number. Sample well data was normalized to aggregated DMSO control wells and plotted versus drug concentration to determine the IC_50_ (infection: blue) and CC_50_ (toxicity: green). Percentage of Control (POC)=(sample well measurement /aggregated DMSO avg)*100 for n=3 replicates. **A)** remdesivir, **B)** entrectinib.

**Table S1.**
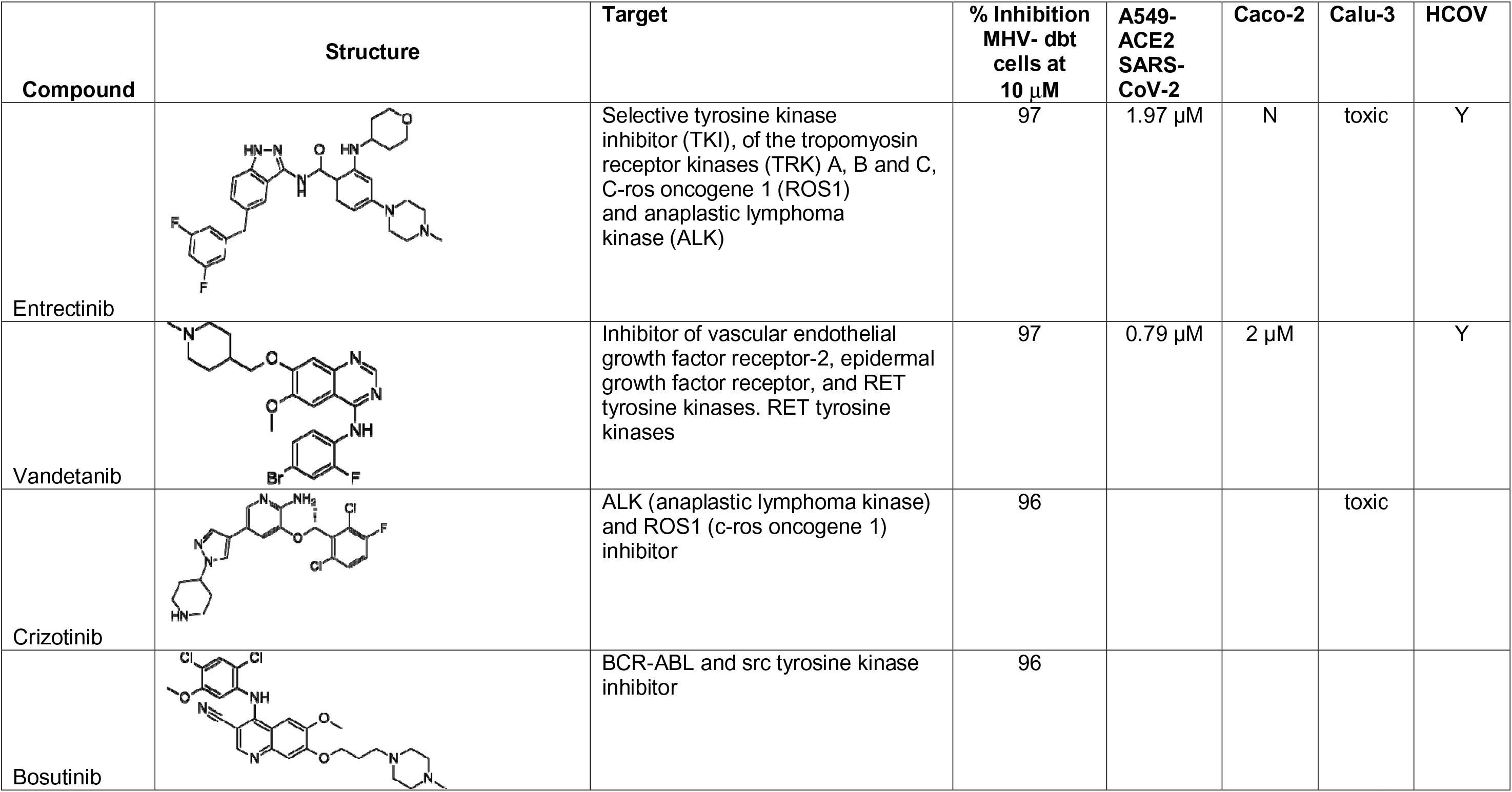
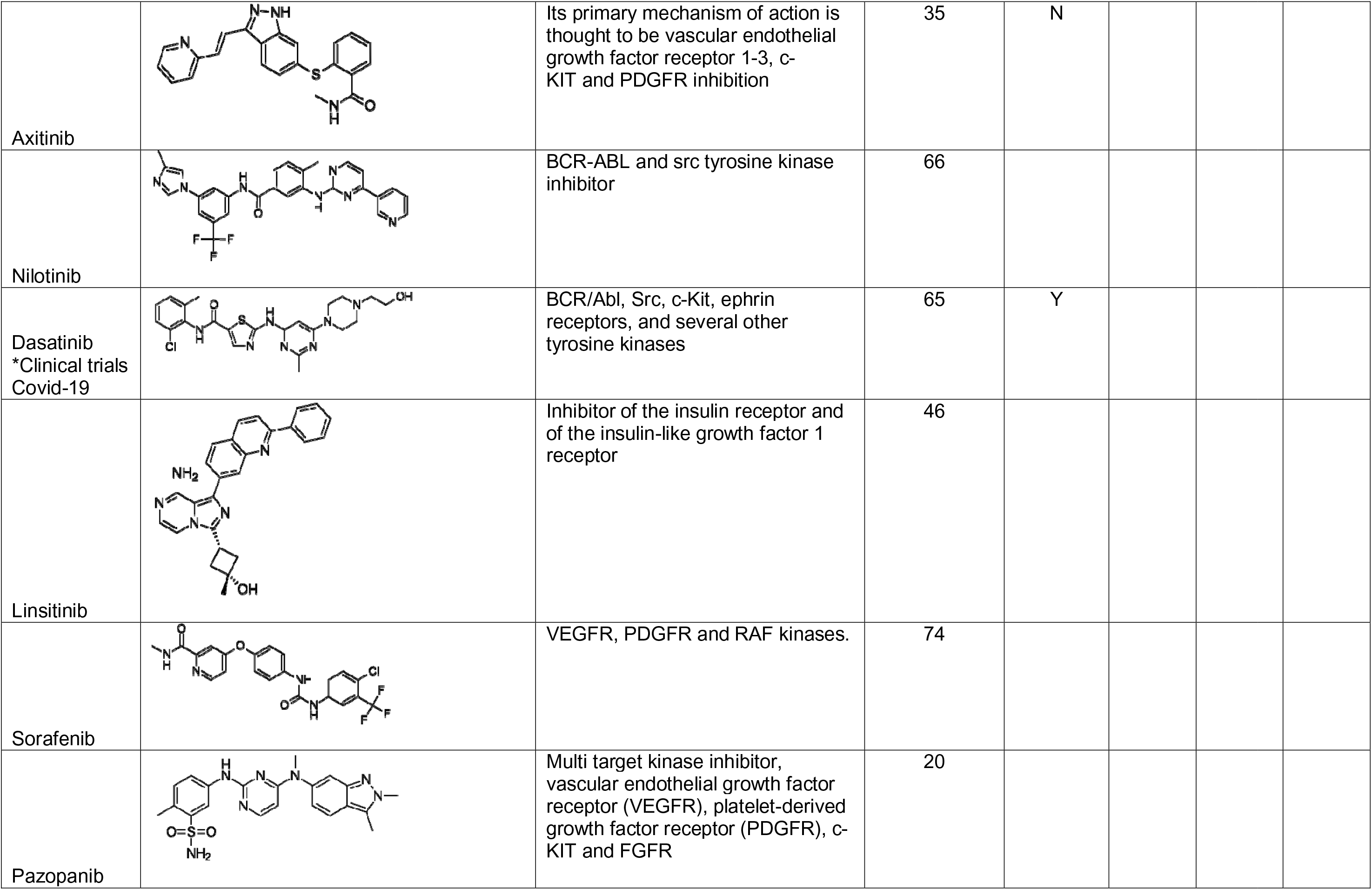
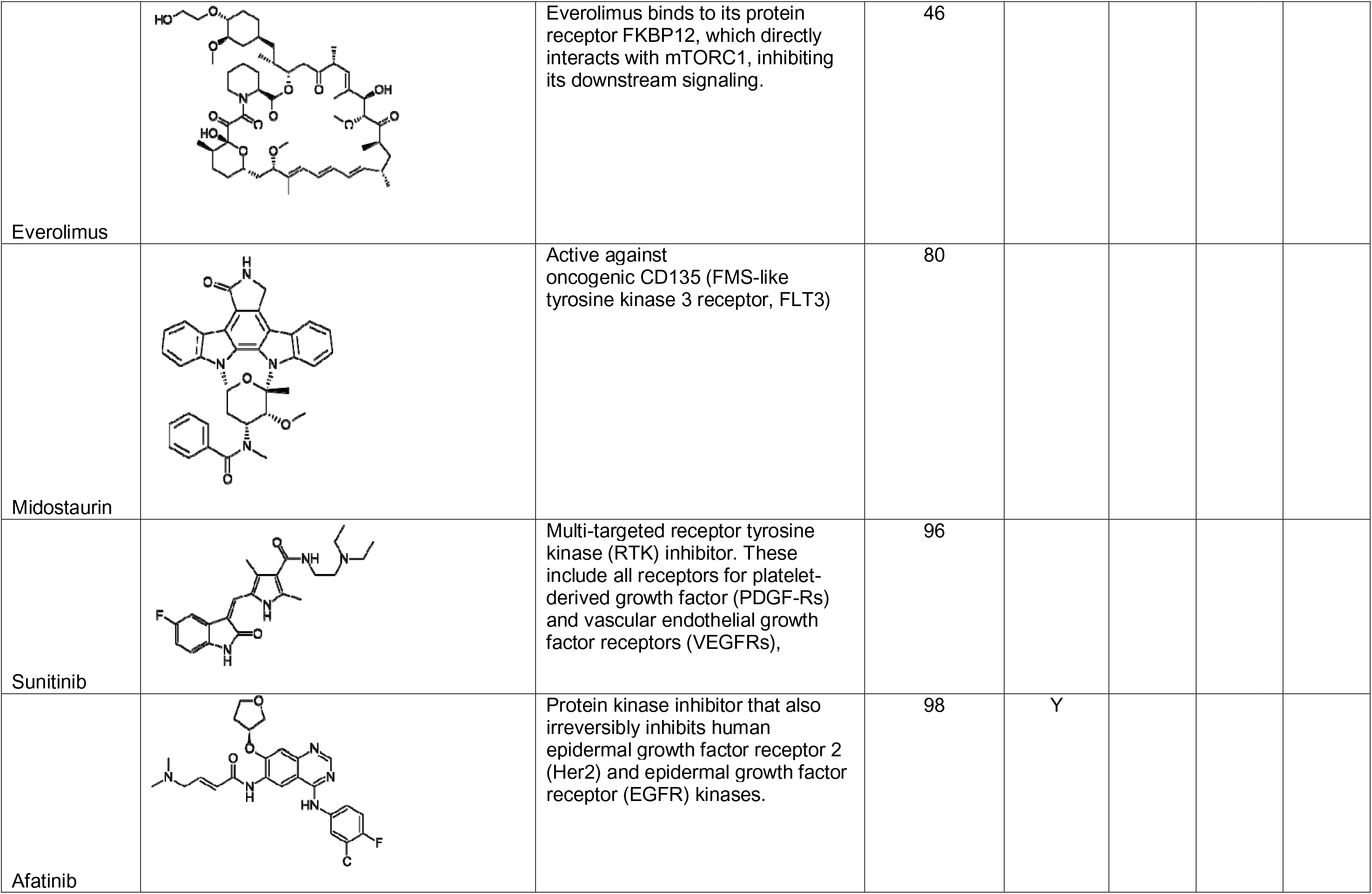
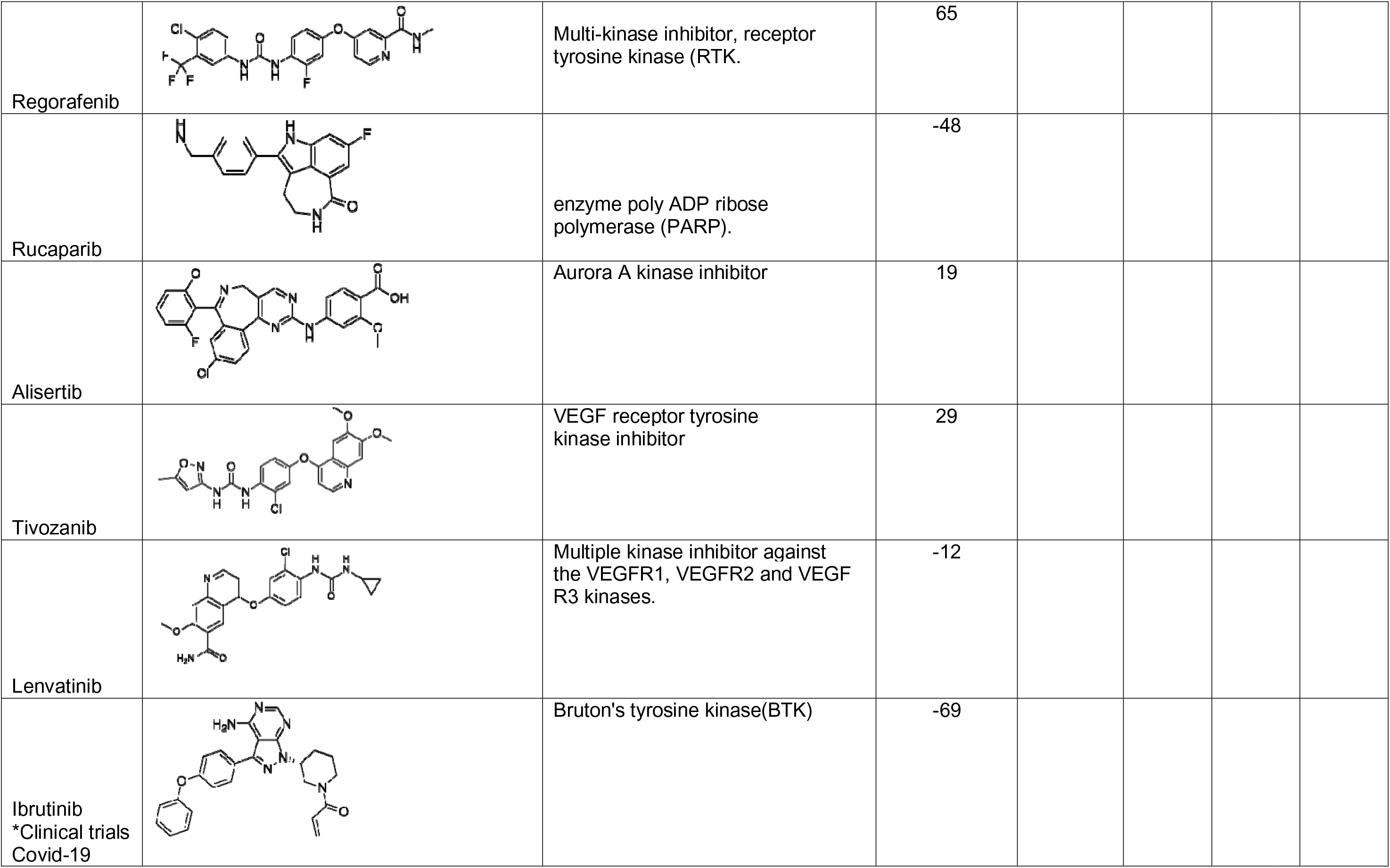
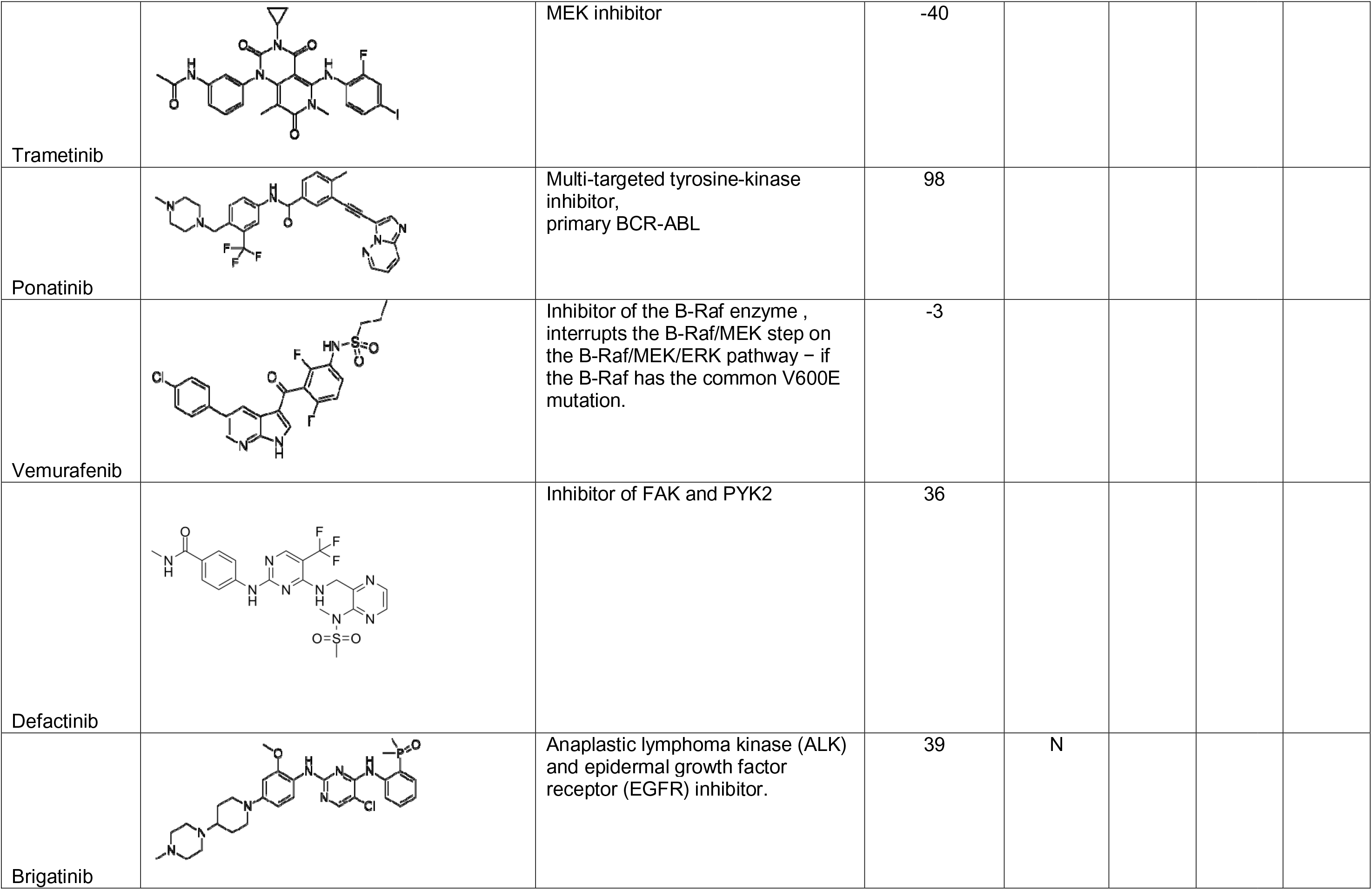
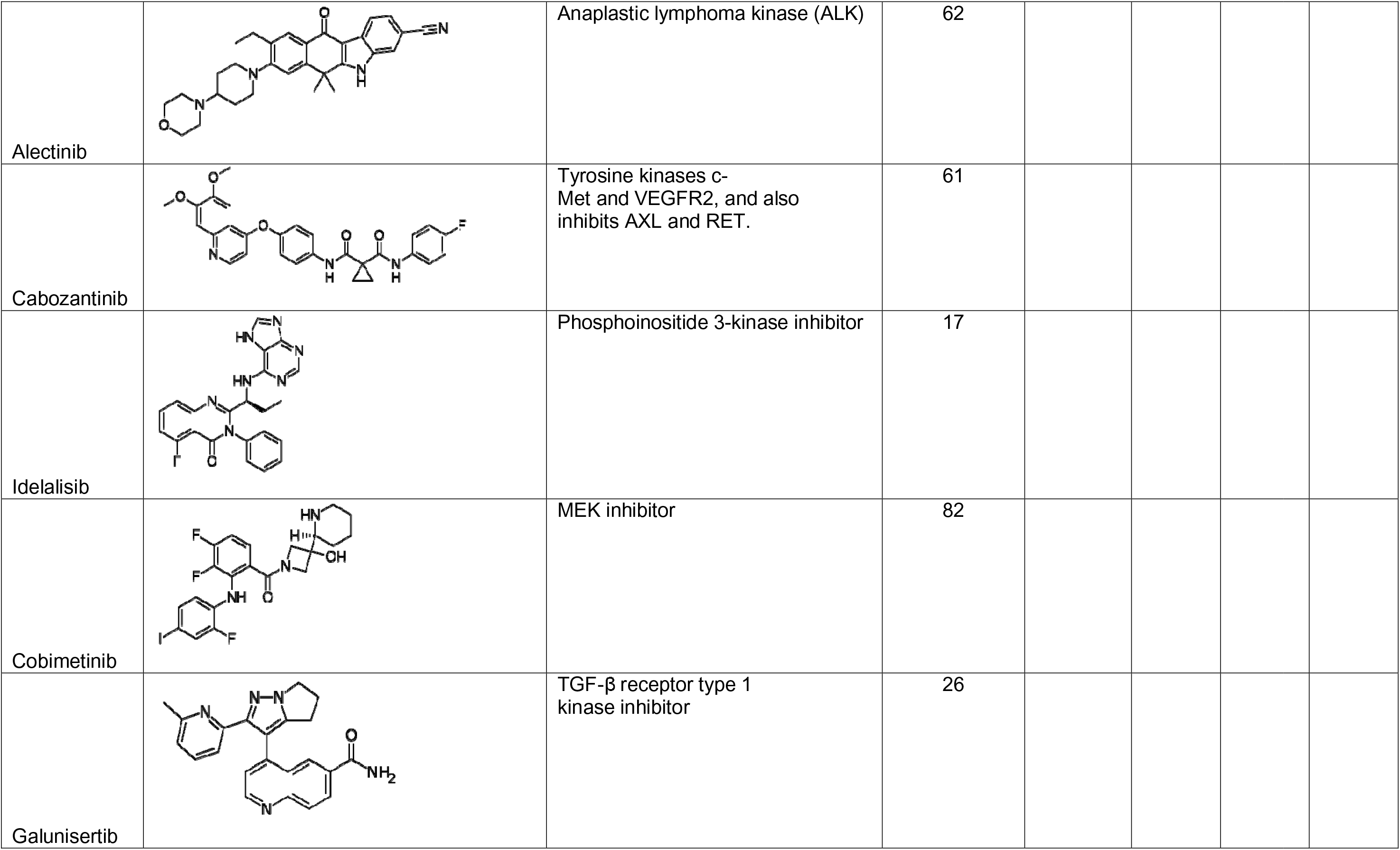
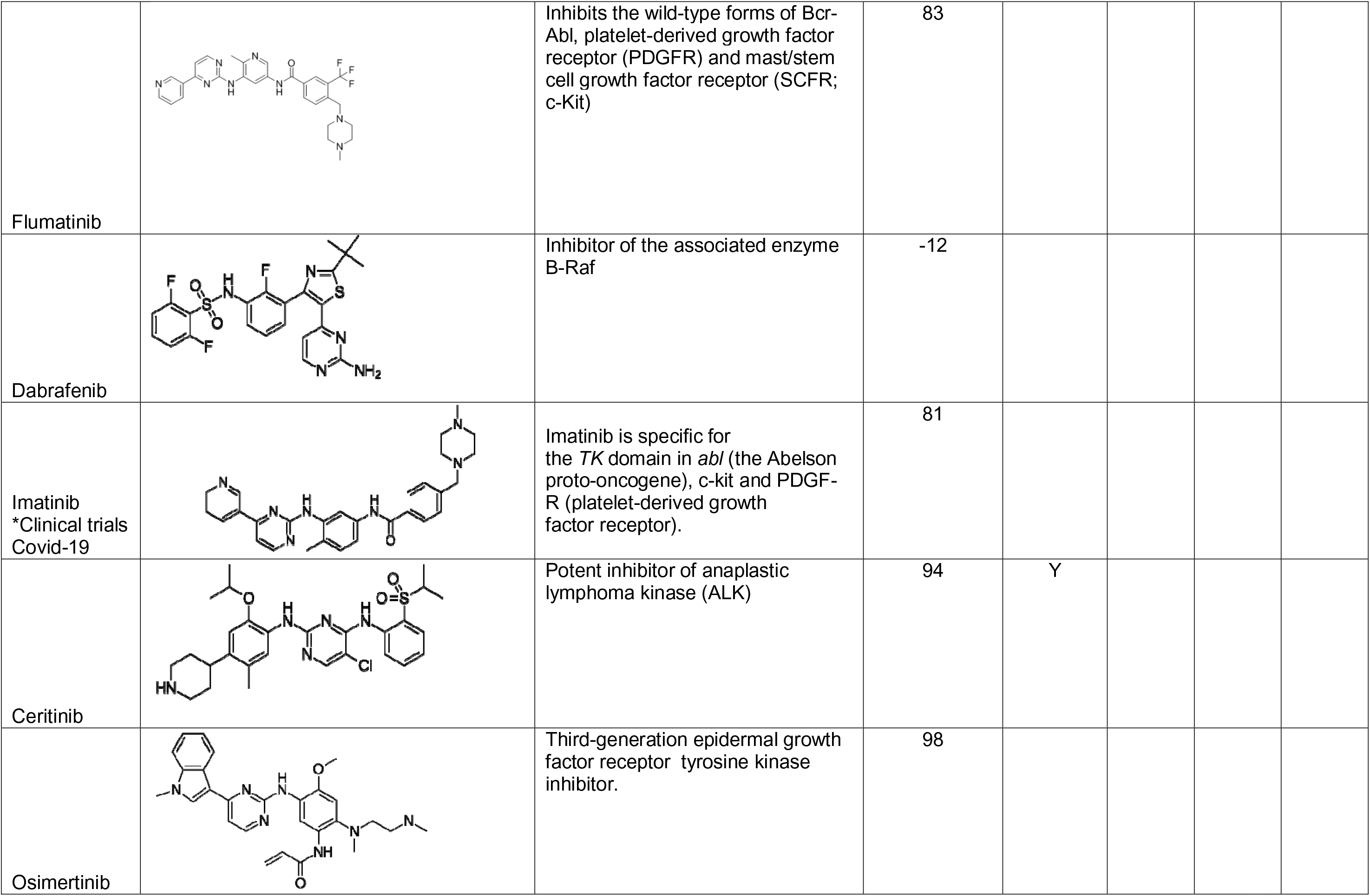
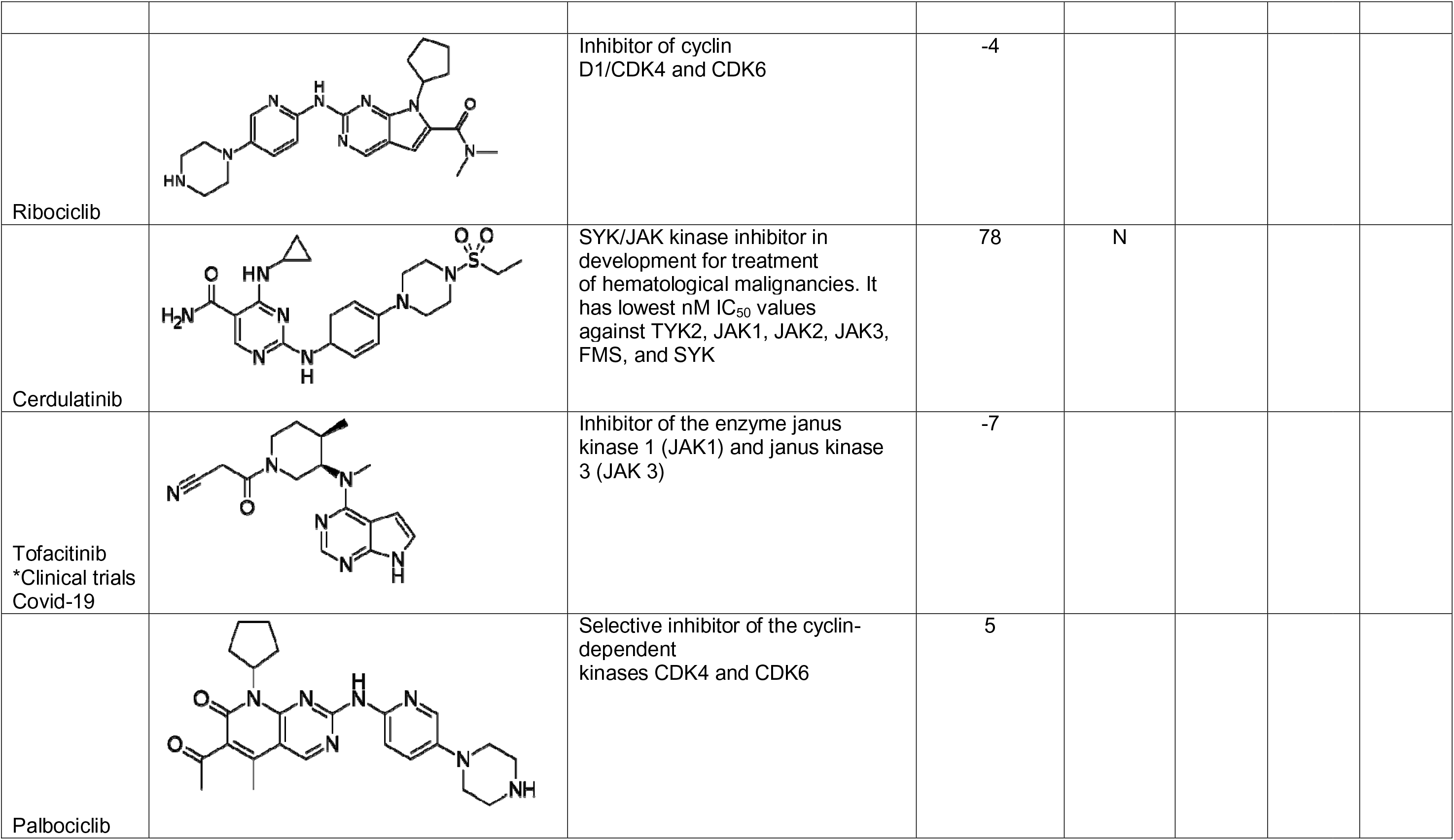
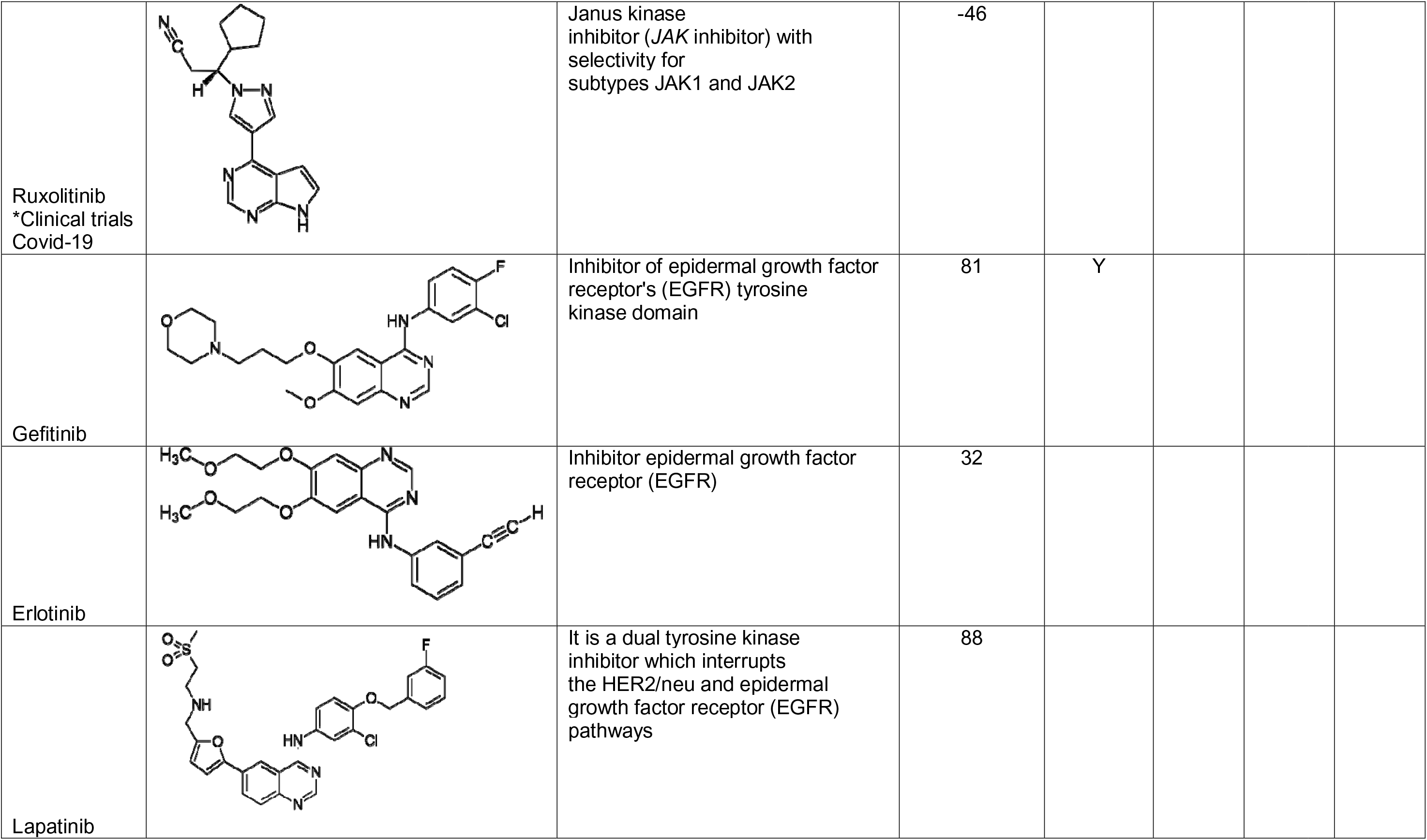
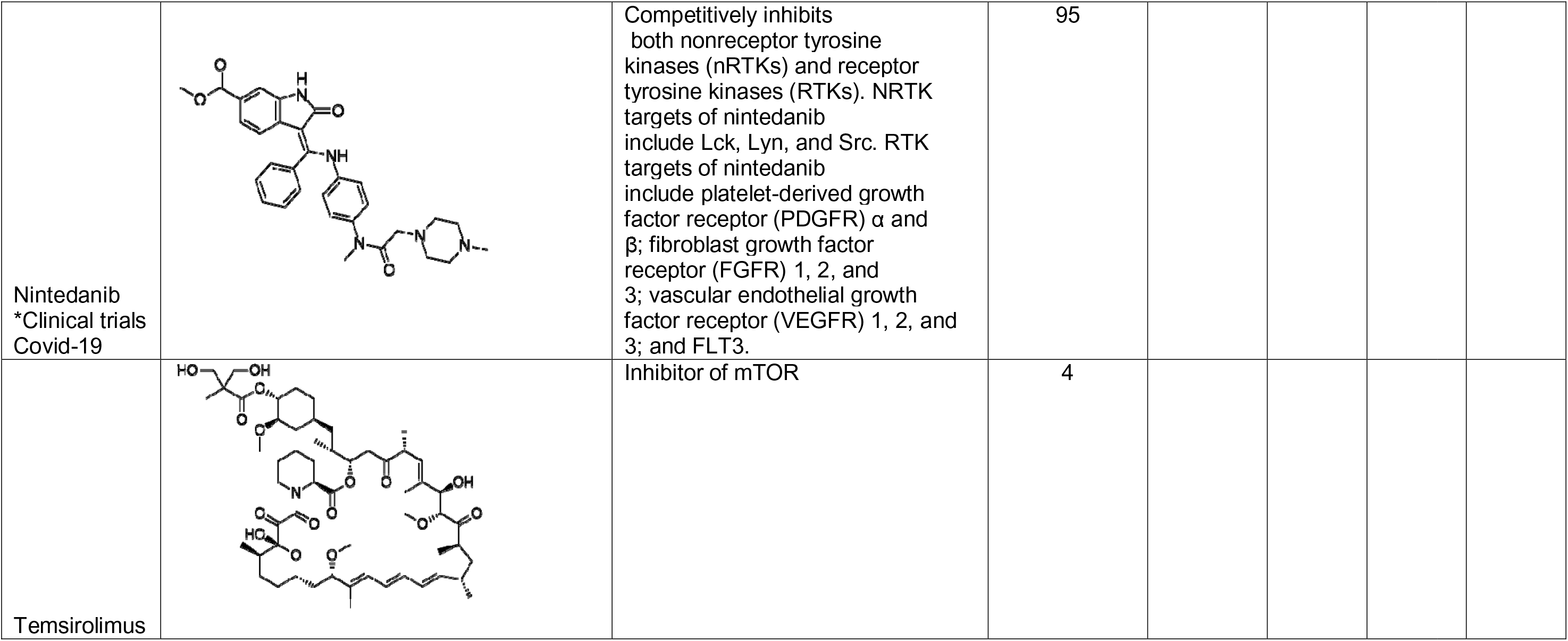
Kinase inhibitors tested against MHV and SARS-CoV-2.

## References

1. Rehman, M.F.U. et al. Novel coronavirus disease (COVID-19) pandemic: A recent mini review. Comput Struct Biotechnol J 19, 612–623 (2021).

2. Kyriakidis, N.C., López-Cortés, A., González, E.V., Grimaldos, A.B. & Prado, E.O. SARS-CoV-2 vaccines strategies: a comprehensive review of phase 3 candidates. NPJ Vaccines 6, 28 (2021).

3. Huang, H.Y. et al. Landscape and progress of global COVID-19 vaccine development. Hum Vaccin Immunother, 1–5 (2021).

4. Eastman, R.T. et al. Remdesivir: A Review of Its Discovery and Development Leading to Emergency Use Authorization for Treatment of COVID-19. ACS Cent Sci 6, 672–683 (2020).

5. Kalil, A.C. et al. Baricitinib plus Remdesivir for Hospitalized Adults with Covid-19. N Engl J Med 384, 795–807 (2021).

6. Horby, P. et al. Dexamethasone in Hospitalized Patients with Covid-19. N Engl J Med 384, 693–704 (2021).

7. Apaydin, C.B., Cinar, G. & Cihan-Ustundag, G. Small-molecule antiviral agents in ongoing clinical trials for COVID-19. Curr Drug Targets (2021).

8. Nalbandian, A. et al. Post-acute COVID-19 syndrome. Nat Med 27, 601–615 (2021).

9. Silva Andrade, B., et al. Long-COVID and Post-COVID Health Complications: An Up-to-Date Review on Clinical Conditions and Their Possible Molecular Mechanisms. Viruses 13 (2021).

10. Ekins, S. et al. Deja vu: Stimulating open drug discovery for SARS-CoV-2. Drug Discov Today 25, 928–941 (2020).

11. Muratov, E.N. et al. A critical overview of computational approaches employed for COVID-19 drug discovery. Chem Soc Rev 50, 9121–9151 (2021).

12. Puhl, A.C. et al. Repurposing the Ebola and Marburg Virus Inhibitors Tilorone, Quinacrine, and Pyronaridine: In Vitro Activity against SARS-CoV-2 and Potential Mechanisms. ACS Omega 6, 7454–7468 (2021).

13. Riva, L. et al. Discovery of SARS-CoV-2 antiviral drugs through large-scale compound repurposing. Nature 586, 113–119 (2020).

14. Amin, S.A., Banerjee, S., Ghosh, K., Gayen, S. & Jha, T. Protease targeted COVID-19 drug discovery and its challenges: Insight into viral main protease (Mpro) and papain-like protease (PLpro) inhibitors. Bioorg Med Chem 29, 115860 (2021).

15. Hoffman, R.L. et al. Discovery of Ketone-Based Covalent Inhibitors of Coronavirus 3CL Proteases for the Potential Therapeutic Treatment of COVID-19. J Med Chem 63, 12725–12747 (2020).

16. Costela-Ruiz, V.J., Illescas-Montes, R., Puerta-Puerta, J.M., Ruiz, C. & Melguizo-Rodríguez, L. SARS-CoV-2 infection: The role of cytokines in COVID-19 disease. Cytokine Growth Factor Rev 54, 62–75 (2020).

17. Chen, R. et al. Cytokine Storm: The Primary Determinant for the Pathophysiological Evolution of COVID-19 Deterioration. Front Immunol 12, 589095 (2021).

18. Roshanravan, N., Seif, F., Ostadrahimi, A., Pouraghaei, M. & Ghaffari, S. Targeting Cytokine Storm to Manage Patients with COVID-19: A Mini-Review. Arch Med Res 51, 608–612 (2020).

19. Coperchini, F. et al. The cytokine storm in COVID-19: Further advances in our understanding the role of specific chemokines involved. Cytokine Growth Factor Rev 58, 82–91 (2021).

20. Felsenstein, S., Herbert, J.A., McNamara, P.S. & Hedrich, C.M. COVID-19: Immunology and treatment options. Clin Immunol 215, 108448 (2020).

21. Peter, A.E., Sandeep, B.V., Rao, B.G. & Kalpana, V.L. Calming the Storm: Natural Immunosuppressants as Adjuvants to Target the Cytokine Storm in COVID-19. Front Pharmacol 11, 583777 (2020).

22. Moradian, N. et al. Cytokine release syndrome: inhibition of pro-inflammatory cytokines as a solution for reducing COVID-19 mortality. Eur Cytokine Netw 31, 81–93 (2020).

23. Beerli, C. et al. Vaccinia virus hijacks EGFR signalling to enhance virus spread through rapid and directed infected cell motility. Nat Microbiol 4, 216–225 (2019).

24. Pleschka, S. et al. Influenza virus propagation is impaired by inhibition of the Raf/MEK/ERK signalling cascade. Nat Cell Biol 3, 301–305 (2001).

25. Roskoski, R. Properties of FDA-approved small molecule protein kinase inhibitors: A 2021 update. Pharmacol Res 165, 105463 (2021).

26. Weisberg, E. et al. Repurposing of Kinase Inhibitors for Treatment of COVID-19. Pharm Res 37, 167 (2020).

27. Baranov, M.V., Bianchi, F. & van den Bogaart, G. The PIKfyve Inhibitor Apilimod: A Double-Edged Sword against COVID-19. Cells 10 (2020).

28. Hoang, T.N. et al. Baricitinib treatment resolves lower-airway macrophage inflammation and neutrophil recruitment in SARS-CoV-2-infected rhesus macaques. Cell 184, 460–475 e421 (2021).

29. Zhang, Q. et al. Heparan sulfate assists SARS-CoV-2 in cell entry and can be targeted by approved drugs in vitro. Cell Discov 6, 80 (2020).

30. Riva, L. et al. Discovery of SARS-CoV-2 antiviral drugs through large-scale compound repurposing. Nature 586, 113–119 (2020).

31. Zhao, H., Mendenhall, M. & Deininger, M.W. Imatinib is not a potent anti-SARS-CoV-2 drug. Leukemia 34, 3085–3087 (2020).

32. Raghuvanshi, R. & Bharate, S.B. Recent Developments in the Use of Kinase Inhibitors for Management of Viral Infections. J Med Chem (2021).

33. Drayman, N. et al. Masitinib is a broad coronavirus 3CL inhibitor that blocks replication of SARS-CoV-2. Science (2021).

34. Corman, V.M. et al. Link of a ubiquitous human coronavirus to dromedary camels. Proc Natl Acad Sci U S A 113, 9864–9869 (2016).

35. Lim, Y.X., Ng, Y.L., Tam, J.P. & Liu, D.X. Human Coronaviruses: A Review of Virus-Host Interactions. Diseases 4 (2016).

36. Fehr, A.R. & Perlman, S. Coronaviruses: an overview of their replication and pathogenesis. Methods Mol Biol 1282, 1–23 (2015).

37. Seidel, S.A. et al. Microscale thermophoresis quantifies biomolecular interactions under previously challenging conditions. Methods 59, 301–315 (2013).

38. Willemsen, M.J. et al. MicroScale Thermophoresis: Interaction analysis and beyond. J Mol Structure 1077, 101–113 (2014).

39. Wrapp, D. et al. Cryo-EM structure of the 2019-nCoV spike in the prefusion conformation. Science 367, 1260–1263 (2020).

40. Chan, K.K. et al. Engineering human ACE2 to optimize binding to the spike protein of SARS coronavirus 2. Science 369, 1261–1265 (2020).

41. Klann, K. et al. Growth Factor Receptor Signaling Inhibition Prevents SARS-CoV-2 Replication. Mol Cell 80, 164–174 e164 (2020).

42. McCray, P.B., Jr. et al. Lethal infection of K18-hACE2 mice infected with severe acute respiratory syndrome coronavirus. J Virol 81, 813–821 (2007).

43. Oladunni, F.S. et al. Lethality of SARS-CoV-2 infection in K18 human angiotensin-converting enzyme 2 transgenic mice. Nat Commun 11, 6122 (2020).

44. Bao, L. et al. The pathogenicity of SARS-CoV-2 in hACE2 transgenic mice. Nature 583, 830–833 (2020).

45. Harvey, W.T. et al. SARS-CoV-2 variants, spike mutations and immune escape. Nat Rev Microbiol 19, 409–424 (2021).

46. Forni, G., Mantovani, A. & COVID-19 Commission of Accademia Nazionale dei Lincei, R.m. COVID-19 vaccines: where we stand and challenges ahead. Cell Death Differ 28, 626–639 (2021).

47. Klann, K. et al. Growth Factor Receptor Signaling Inhibition Prevents SARS-CoV-2 Replication. Mol Cell 80, 164–174.e164 (2020).

48. Frampton, J.E. Vandetanib: in medullary thyroid cancer. Drugs 72, 1423–1436 (2012).

49. Wang, J. et al. Specific cytokines in the inflammatory cytokine storm of patients with COVID-19-associated acute respiratory distress syndrome and extrapulmonary multiple-organ dysfunction. Virol J 18, 117 (2021).

50. Aid, M. et al. Vascular Disease and Thrombosis in SARS-CoV-2-Infected Rhesus Macaques. Cell 183, 1354–1366 e1313 (2020).

51. Huang, C. et al. Clinical features of patients infected with 2019 novel coronavirus in Wuhan, China. Lancet 395, 497–506 (2020).

52. Herold, T. et al. Elevated levels of IL-6 and CRP predict the need for mechanical ventilation in COVID-19. J Allergy Clin Immunol 146, 128–136.e124 (2020).

53. Zhang, Q. et al. Inborn errors of type I IFN immunity in patients with life-threatening COVID-19. Science 370 (2020).

54. Xia, H. et al. Evasion of Type I Interferon by SARS-CoV-2. Cell Rep 33, 108234 (2020).

55. Yuen, C.K. et al. SARS-CoV-2 nsp13, nsp14, nsp15 and orf6 function as potent interferon antagonists. Emerg Microbes Infect 9, 1418–1428 (2020).

56. Molony, R.D. et al. Aging impairs both primary and secondary RIG-I signaling for interferon induction in human monocytes. Sci Signal 10 (2017).

57. Hu, R., Xia, C.Q., Butfiloski, E. & Clare-Salzler, M. Effect of high glucose on cytokine production by human peripheral blood immune cells and type I interferon signaling in monocytes: Implications for the role of hyperglycemia in the diabetes inflammatory process and host defense against infection. Clin Immunol 195, 139–148 (2018).

58. Chi, Y. et al. Serum Cytokine and Chemokine Profile in Relation to the Severity of Coronavirus Disease 2019 in China. J Infect Dis 222, 746–754 (2020).

59. Abers, M.S., et al. An immune-based biomarker signature is associated with mortality in COVID-19 patients. JCI Insight 6 (2021).

60. Sierra, B. et al. Association of Early Nasopharyngeal Immune Markers With COVID-19 Clinical Outcome: Predictive Value of CCL2/MCP-1. Open Forum Infect Dis 7, ofaa407 (2020).

61. Kaner, R.J. et al. Lung overexpression of the vascular endothelial growth factor gene induces pulmonary edema. Am J Respir Cell Mol Biol 22, 657–664 (2000).

62. Marti, H.H. & Risau, W. Systemic hypoxia changes the organ-specific distribution of vascular endothelial growth factor and its receptors. Proc Natl Acad Sci U S A 95, 15809–15814 (1998).

63. Lee, C.G. et al. Vascular endothelial growth factor (VEGF) induces remodeling and enhances TH2-mediated sensitization and inflammation in the lung. Nat Med 10, 1095–1103 (2004).

64. Pang, J. et al. Efficacy and tolerability of bevacizumab in patients with severe Covid-19. Nat Commun 12, 814 (2021).

65. Weisberg, E. et al. Repurposing of Kinase Inhibitors for Treatment of COVID-19. Pharm Res 37, 167 (2020).

66. Sheahan, T.P. et al. Broad-spectrum antiviral GS-5734 inhibits both epidemic and zoonotic coronaviruses. Sci Transl Med 9 (2017).

67. Beigel, J.H. et al. Remdesivir for the Treatment of Covid-19 - Final Report. N Engl J Med 383, 1813–1826 (2020).

68. Group, R.C. et al. Dexamethasone in Hospitalized Patients with Covid-19. N Engl J Med 384, 693–704 (2021).

69. Guimaraes, P.O. et al. Tofacitinib in Patients Hospitalized with Covid-19 Pneumonia. N Engl J Med 385, 406–415 (2021).

70. Kyriazopoulou, E. et al. Author Correction: Early treatment of COVID-19 with anakinra guided by soluble urokinase plasminogen receptor plasma levels: a double-blind, randomized controlled phase 3 trial. Nat Med 27, 1850 (2021).

71. Drayman, N. et al. Masitinib is a broad coronavirus 3CL inhibitor that blocks replication of SARS-CoV-2. Science 373, 931–936 (2021).

72. Martin, P. et al. Pharmacokinetics of vandetanib: three phase I studies in healthy subjects. Clin Ther 34, 221–237 (2012).

73. Jeong, W., Doroshow, J.H. & Kummar, S. United States Food and Drug Administration approved oral kinase inhibitors for the treatment of malignancies. Curr Probl Cancer 37, 110–144 (2013).

74. Prete, A. et al. Thyroid cancer and COVID-19: experience at one single thyroid disease referral center. Endocrine 72, 332–339 (2021).

## Supplemental References

1. Premkumar, L., et al. The receptor binding domain of the viral spike protein is an immunodominant and highly specific target of antibodies in SARS-CoV-2 patients. Sci Immunol 5 (2020).

2. Puhl, A.C. et al. Repurposing the Ebola and Marburg Virus Inhibitors Tilorone, Quinacrine, and Pyronaridine:. ACS Omega 6, 7454–7468 (2021).

3. Hou, Y.J. et al. SARS-CoV-2 Reverse Genetics Reveals a Variable Infection Gradient in the Respiratory Tract. Cell 182, 429–446 e414 (2020).

4. Yount, B., Denison, M.R., Weiss, S.R. & Baric, R.S. Systematic assembly of a full-length infectious cDNA of mouse hepatitis virus strain A59. J Virol 76, 11065–11078 (2002).

5. Spearman, C. The method of ‘right and wrong cases’ (‘constant stimuli’) without Gauss’s formulae. Brit J Psychol 2, 227–242 (1908).

6. Kärber, G. Beitrag zur kollektiven behandlung pharmakologischer reihenversuche. Archiv f Experiment Pathol u Pharmakol 162, 480–483 (1931).

7. McCray, P.B. et al. Lethal infection of K18-hACE2 mice infected with severe acute respiratory syndrome coronavirus. J Virol 81, 813–821 (2007).

8. Oladunni, F.S. et al. Lethality of SARS-CoV-2 infection in K18 human angiotensin-converting enzyme 2 transgenic mice. Nat Commun 11, 6122 (2020).

9. Bao, L. et al. The pathogenicity of SARS-CoV-2 in hACE2 transgenic mice. Nature 583, 830–833 (2020).

10. Yinda, C.K. et al. K18-hACE2 mice develop respiratory disease resembling severe COVID-19. PLoS Pathog 17, e1009195 (2021).

11. Arce, V.M. & Costoya, J.A. SARS-CoV-2 infection in K18-ACE2 transgenic mice replicates human pulmonary disease in COVID-19. Cell Mol Immunol 18, 513–514 (2021).

12. Moreau, G.B. et al. Evaluation of K18-. Am J Trop Med Hyg 103, 1215–1219 (2020).

13. Winkler, E.S. et al. Publisher Correction: SARS-CoV-2 infection of human ACE2-transgenic mice causes severe lung inflammation and impaired function. Nat Immunol 21, 1470 (2020).

14. Zheng, J. et al. COVID-19 treatments and pathogenesis including anosmia in K18-hACE2 mice. Nature 589, 603–607 (2021).

15. Hsia, C.C., Hyde, D.M., Ochs, M., Weibel, E.R. & Structure, A.E.J.T.F.o.Q.A.o.L. An official research policy statement of the American Thoracic Society/European Respiratory Society: standards for quantitative assessment of lung structure. Am J Respir Crit Care Med 181, 394–418 (2010).

